# Genes and pathways implicated in tetralogy of Fallot revealed by ultra-rare variant burden analysis in 231 genome sequences

**DOI:** 10.1101/2020.03.02.972653

**Authors:** Roozbeh Manshaei, Daniele Merico, Miriam S. Reuter, Worrawat Engchuan, Bahareh A. Mojarad, Rajiv Chaturvedi, Tracy Heung, Giovanna Pellecchia, Mehdi Zarrei, Thomas Nalpathamkalam, Reem Khan, John B. A. Okello, Eriskay Liston, Meredith Curtis, Ryan K.C. Yuen, Christian R. Marshall, Rebekah K. Jobling, Stephen W. Scherer, Raymond H. Kim, Anne S. Bassett

## Abstract

Recent genome-wide studies of rare genetic variants have begun to implicate novel mechanisms for tetralogy of Fallot (TOF), a severe congenital heart defect (CHD).

To provide statistical support for case-only data without parental genomes, we re-analyzed genome sequences of 231 individuals with TOF or related CHD. We adapted a burden test originally developed for *de novo* variants to assess singleton variant burden in individual genes, and in gene-sets corresponding to functional pathways and mouse phenotypes, accounting for highly correlated gene-sets, and for multiple testing.

The gene burden test identified a significant burden of deleterious missense variants in *NOTCH1* (Bonferroni-corrected p-value <0.01). These *NOTCH1* variants showed significant enrichment for those affecting the extracellular domain, and especially for disruption of cysteine residues forming disulfide bonds (OR 39.8 vs gnomAD). Individuals with *NOTCH1* variants, all with TOF, were enriched for positive family history of CHD. Other genes not previously implicated in TOF had more modest statistical support and singleton missense variant results were non-significant for gene-set burden. For singleton truncating variants, the gene burden test confirmed significant burden in *FLT4.* Gene-set burden tests identified a cluster of pathways corresponding to VEGF signaling (*FDR*=0%), and of mouse phenotypes corresponding to abnormal vasculature (*FDR*=0.8%), that suggested additional candidate genes not previously identified (e.g., *WNT5A* and *ZFAND5*). Analyses using unrelated sequencing datasets supported specificity of the findings for CHD.

The findings support the importance of ultra-rare variants disrupting genes involved in VEGF and NOTCH signaling in the genetic architecture of TOF. These proof-of-principle data indicate that this statistical methodology could assist in analyzing case-only sequencing data in which ultra-rare variants, whether *de novo* or inherited, contribute to the genetic etiopathogenesis of a complex disorder.

**Author summary:** We analyzed the ultra-rare nonsynonymous variant burden for genome sequencing data from 231 individuals with congenital heart defects, most with tetralogy of Fallot. We adapted a burden test originally developed for *de novo* variants. In line with other studies, we identified a significant truncating variant burden for *FLT4* and deleterious missense burden for *NOTCH1*, both passing a stringent Bonferroni multiple-test correction. For *NOTCH1*, we observed frequent disruption of cysteine residues establishing disulfide bonds in the extracellular domain. We also identified genes with BH-FDR <10% that were not previously implicated. To overcome limited power for individual genes, we tested gene-sets corresponding to functional pathways and mouse phenotypes. Gene-set burden of truncating variants was significant for vascular endothelial growth factor signaling and abnormal vasculature phenotypes. These results confirmed previous findings and suggested additional candidate genes for experimental validation in future studies. This methodology can be extended to other case-only sequencing data in which ultra-rare variants make a substantial contribution to genetic etiology.

## Introduction

Congenital heart defects (CHD) occur in 8/1000 live births and are a leading cause of mortality from birth defects (1), with a wide spectrum of severity (2). Among CHD, tetralogy of Fallot (TOF) is the most common of the more severe (cyanotic) conditions. Individuals with TOF present with a combination of abnormalities (pulmonary valve stenosis, right ventricular hypertrophy, ventricular septal defect and overriding aorta) that together lead to insufficient tissue oxygenation.

Genetic factors are major contributors to the etiology of TOF. These include 20% of patients with pathogenic copy number variants (CNV) or larger chromosomal anomalies (3,4). Recent studies have also begun to elucidate the genome-wide role of rare variants at the sequence level, including substitutions and small insertions/deletions.

In a multi-centre exome sequencing study of various CHD that focused on loss-of-function variants and included parental sequencing data enabling *de novo* variant identification, the TOF sub-group drove a significant genome-wide burden finding (p-value ≤1.3×10^−6^) of *de novo* and rare inherited heterozygous truncating variants for a novel gene, *FLT4* (5). Of 9 probands with *FLT4* truncating variants, corresponding to 2.3% of the TOF group, 7 were inherited with evidence of incomplete penetrance (5).

In an independent case-only study using whole genome sequencing (WGS), we investigated 175 adults with TOF for rare loss-of-function variants (including structural variants) disrupting *FLT4* and other vascular endothelial growth factor (VEGF) pathway genes predicted to be haploinsufficient based on the ExAC pLI index (6,7). We identified seven truncating variants in *FLT4*, two in *KDR*, and one each in *BCAR1*, *FGD5*, *FOXO1*, *IQGAP1* and *PRDM1*, corresponding in aggregate to 8% of participants; all variants were absent from public databases^6^. The results suggested the importance of VEGF signaling; however, the statistical burden was not systematically investigated. Another recent multi-centre exome sequencing study of 829 patients with TOF reported genome-wide significant (p-value ≤5×10^−8^) excess of ultra-rare (absent from a public exome database and other reference data) deleterious variants for *FLT4* and *NOTCH1* (8). Loss-of-function variants predominated for *FLT4*, and missense variants for *NOTCH1* (8).

In this study, we undertook a comprehensive statistical re-analysis of the cohort with WGS data previously investigated for ultra-rare variants in the VEGF pathway (6). In an attempt to boost power, we included the sequencing data available for 56 CHD cases as well as for the original 175 TOF cases (n=231 total). We focused on ultra-rare truncating (stop-gain, frameshift and splice site altering) and missense variants that were not reported in the gnomAD database and were identified in only one proband, i.e., that were singletons. We tested burden by adapting a test originally developed for *de novo* variants by rescaling the mutation probability for singletons. Since singletons are enriched in *de novo* variants and are likely to have arisen recently, this is an appropriate extension of the test. To boost power, we additionally tested gene-sets corresponding to (a) functional pathways, derived from Gene Ontology (GO) (9), BioCarta (http://cgap.nci.nih.gov/Pathways/BioCarta_Pathways/), Kyoto Encyclopedia of Genes and Genomes (KEGG) (10) (http://www.genome.jp/kegg/), REACTOME (11), NCI-Nature Pathway Interaction Database (PID) (http://pid.nci.nih.gov); and (b) phenotypes in mouse orthologues, derived from Mouse Genome Informatics (MGI) and based on the Mouse Phenotype Ontology (MPO) classification (12). To control for correlations between highly overlapping gene-sets that could lead to incorrect multiple p-value corrections, we adopted a greedy step-down approach to cluster gene-sets with highly overlapping genes. A sampling-based false discovery rate (FDR) was then estimated. We did not analyze structural variants because no broadly-accepted probabilistic framework has yet been developed to determine the statistical significance of their burden.

## Results

### Identification of singleton variants

Variant calls from the CHD WGS data-set were filtered to retain only high-quality singletons, that were then categorized as truncating or missense based on their effect on the principal transcript (see Materials and Methods for details). With respect to the 2,003 truncating singleton variants initially identified, 868 variants remained after applying the low quality and frameshift indel filter, 764 after applying the principal transcript effect filter, 752 after applying the splice site alteration filter, and finally 642 after considering a maximum of one singleton variant per gene per subject. For the 4,324 missense variants initially identified, 3,521 remained after applying the low-quality filter, 3,359 after applying the principal transcript filter, and finally, 3,293 singleton missense variants after considering a maximum of one singleton variant per gene per subject. We then tested these ultra-rare truncating and missense singleton variants for gene and gene-set burden (see Figure 1 for an overview of the analysis workflow; all ultra-rare singleton variants identified are listed in Supplementary Table S1). For all analyses we tested truncating and missense variants separately because of the likely differences in the genetic architecture of these variant types.

**Figure 1).**
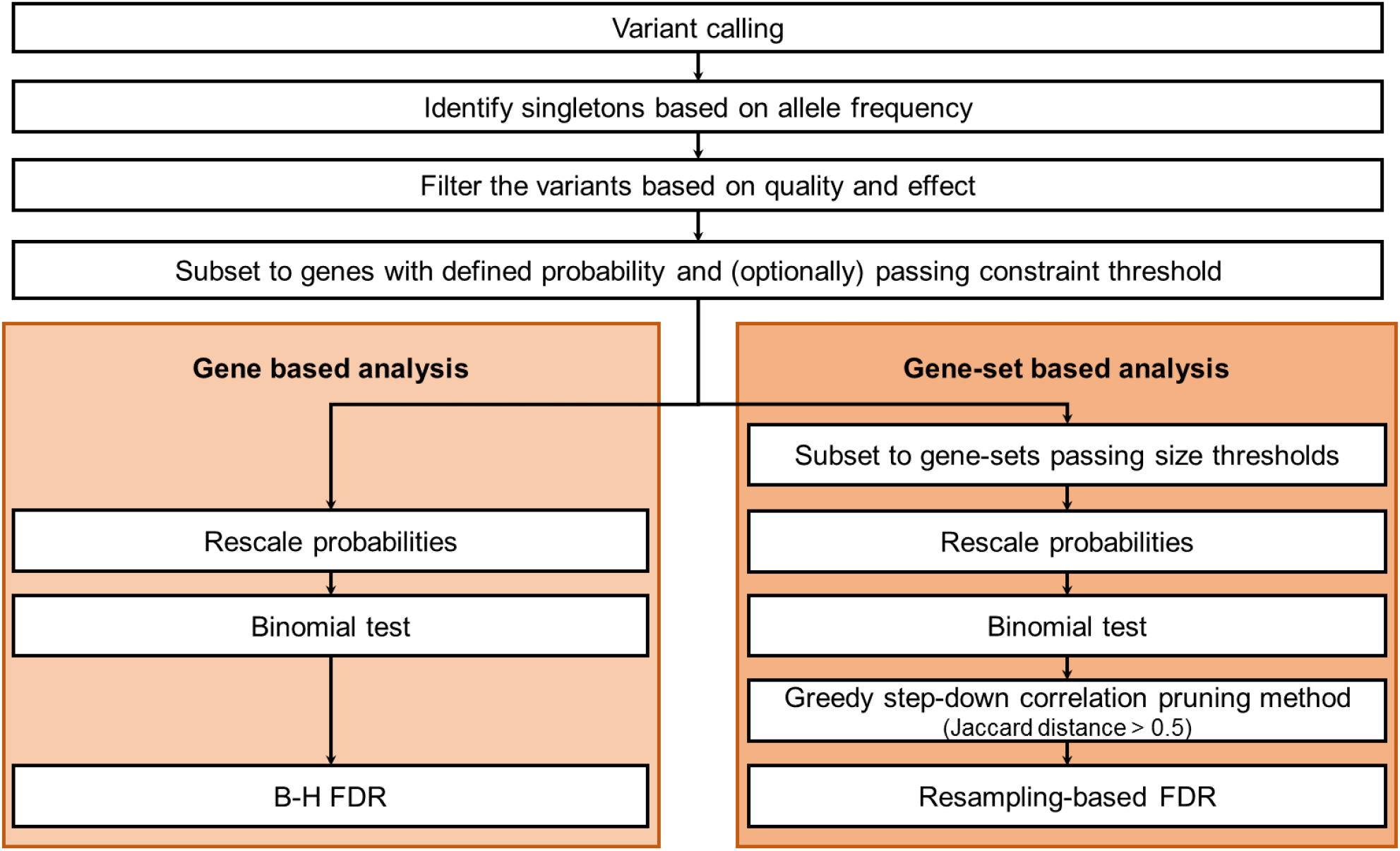
General gene and gene-set burden analyses overview.

### Gene burden results

Genes were tested separately for the burden of singleton truncating and missense variants, using a binomial test based on rescaled *de novo* mutation probabilities (as described in the Materials and Methods). We performed multiple test correction on all genes with a defined probability, and also on a more constrained subset: for truncating variants, gnomAD LOF o/e < 0.35; for missense variants, gnomAD missense o/e < 0.75 (where *o/e* indicates observed/expected; see Supplementary Figure S1 for the relation to the *pLI* and missense *z-score* constraint indexes). Constrained genes are presumed to be more likely to contribute to disease, since they are under negative selection; these thresholds were specifically set to include moderately constrained genes, considering the incomplete penetrance observed for TOF (5,6,8). There were 603 genes with at least one truncating singleton variant, of which 163 passed the constraint threshold; there were 2801 genes with at least one singleton missense variant, of which 739 passed the constraint threshold (see Supplementary Table S2 and Supplementary Table S3 for details). To assess the validity of the gene burden results, we performed several additional analyses: (a) we checked the distribution of observed p-values compared to expected p-values, to monitor for systematic p-value inflation; (b) we compared the p-values obtained for CHD to those obtained for WGS data available for 263 individuals with schizophrenia, processed in exactly the same way; (c) we reassessed the burden by comparing to gnomAD singletons.

For truncating variants, there was only one constrained gene (of the 163 with at least one singleton truncating variant) with significant burden: *FLT4* (uncorrected p-value = 9.56×10^−12^, BH-FDR = 6.99×10^−8^, Bonferroni-corrected p-value = 6.99×10^−8^). When testing all genes (including 603 with at least one singleton truncating variant), in addition to *FLT4*, we identified *CLDN9* as significant at an FDR threshold of 10% (uncorrected p-value = 7.80×10^−6^, BH-FDR = 0.073, Bonferroni-corrected p-value = 0.145) (see Table 1 and Supplementary Table S2 for all details). There was no evidence of genome-wide inflation in either analysis (see Figure 2 and Supplementary Figure S4).

**Table 1).**
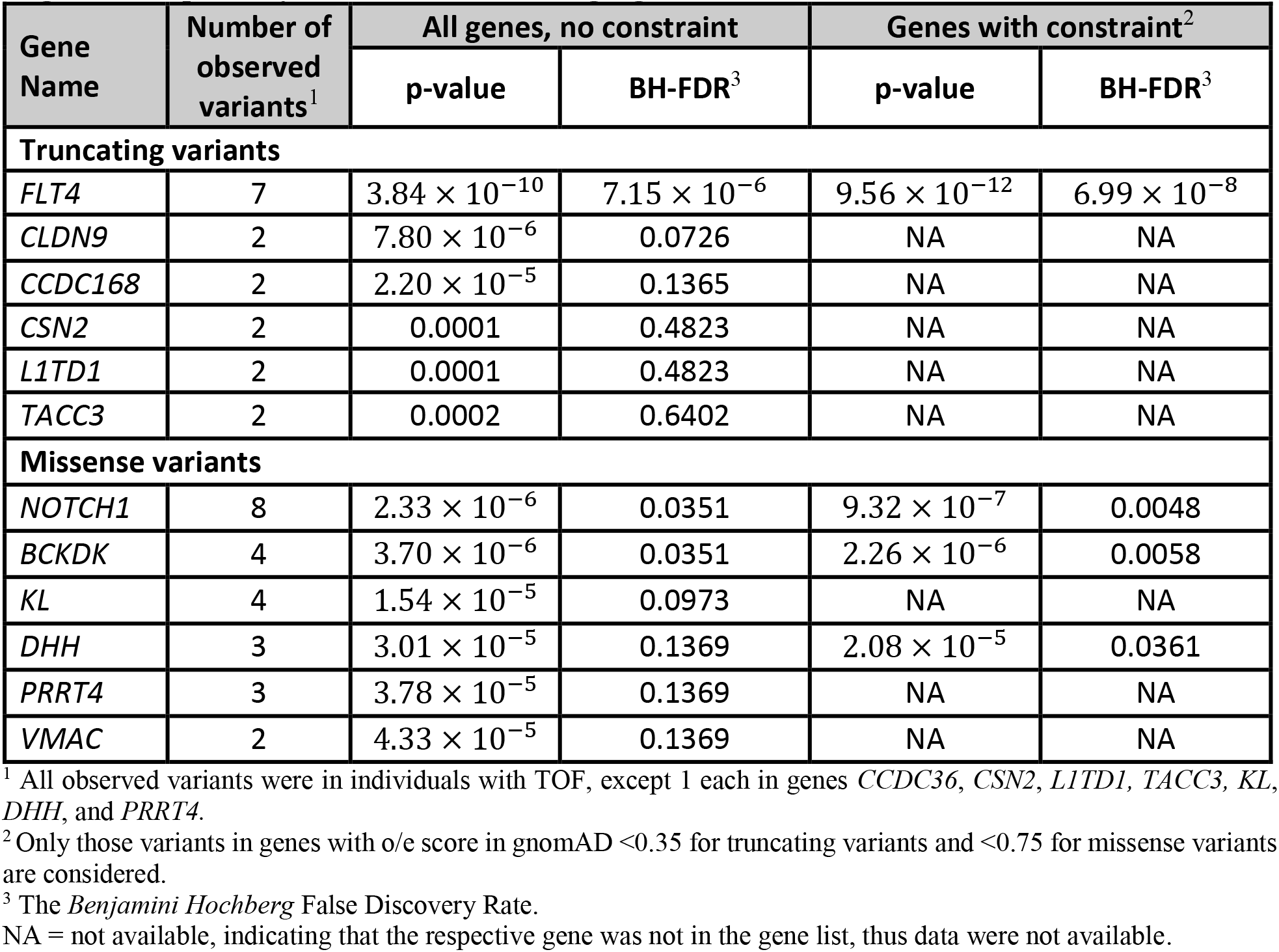
Top six nominally significant genes with ultra-rare (singleton) variants identified in 231 individuals with CHD, as inferred from gene-based burden analyses for truncating and missense singletons, respectively, with and without using a gene constraint cut-off.

**Figure 2).**
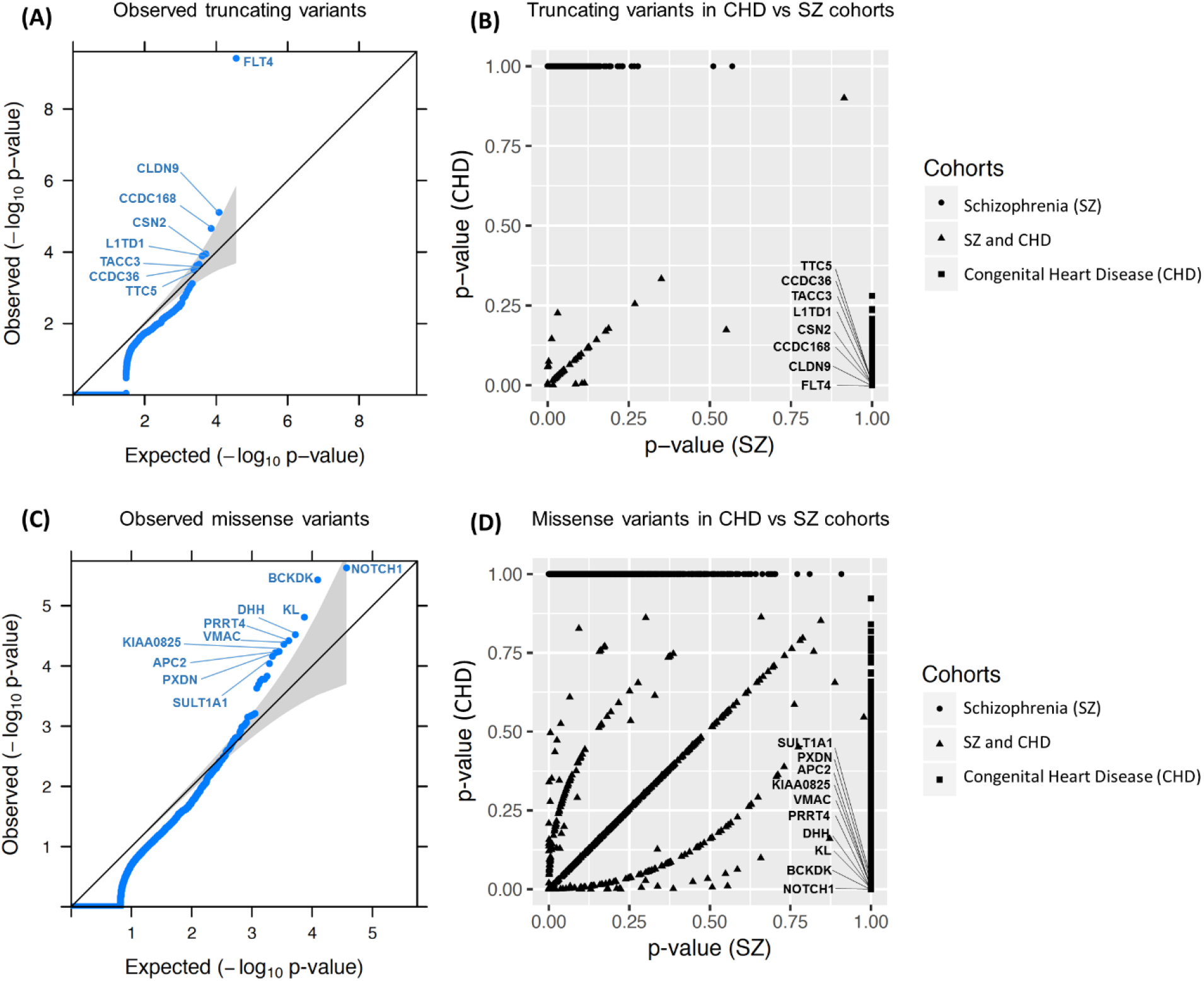
Gene burden analysis results for all genes. (A) and (C) show the quantile-quantile (QQ) plots obtained for all ultra-rare truncating and missense variants in CHD, respectively (i.e., not setting any gene constraint cutoff). The QQ plots represent the scatter plots of the −log10(p-value) expected under the null hypothesis of no genetic association versus the observed −log10(p-value) for all 231 CHD samples. Grey shading indicates the 95% confidence interval. (B) and (D) represent scatter plots of gene burden p-values for truncating and missense variants respectively, comparing the CHD and schizophrenia WGS data. Names of the top 8 and 10 genes identified for truncating (A) and missense (C) variants, respectively, are shown (results for top 6 of each are presented in Table 1); *FLT4* (A) and *NOTCH1* (C) were the most significant genes identified, neither with any observation in the comparison schizophrenia cohort (B, D). These plots were generated based on the genes without constraint on o/e score. Supplementary Figure S4 shows results for genes with constraint.

Considering the top-associated CHD genes without using a constraint threshold, none had a similar p-value in the schizophrenia sequencing data. When applying the constraint threshold, a single top-associated gene that failed the 10% BH-FDR threshold (*ATXN3*) appeared to have a somewhat similar p-value for schizophrenia. However, visualization of the bam files for the schizophrenia data re-classified those variants to be in-frame polymorphisms (see Supplementary Table S4). For *FLT4* and *CLDN9,* where BH-FDR was under the 10% threshold, we evaluated the truncating singleton burden in CHD compared to that in gnomAD: *FLT4* had an even more significant association (uncorrected p-value = 2.43×10^−15^, BH-FDR = 4.01×10^−11^), whereas *CLDN9* was less significant (uncorrected p-value = 7.8×10^−4^, BH-FDR =1), leading us to question the validity of its association to CHD (see Supplementary Table S5 and Supplementary Figure S2). Restricting to constrained genes may have some utility in prioritizing genes, but these results are too limited to draw robust general conclusions.

Of the 739 genes with singleton missense variants that passed the constraint threshold, there were three genes that passed the 10% FDR threshold: *NOTCH1* (uncorrected p-value = 9.32×10^−7^, BH-FDR=0.0048, Bonferroni-corrected p-value = 4.85×10^−3^), *BCKDK* (uncorrected p-value = 2.26×10^− 6^, BH-FDR=0.0058, Bonferroni-corrected p-value = 0.018), and *DHH* (uncorrected p-value = 2.08×10^−5^, BH-FDR=0.0361, Bonferroni-corrected p-value = 0.108); see Table 1 and Supplementary Table S3 for further details. When considering all 2801 genes with singleton missense variants, regardless of constraint, the BH-FDR for gene *DHH* (0.1369) was less significant, but another gene, *KL*, passed the 10% BH-FDR cut-off (uncorrected p-value = 1.54×10^− 5^, BH-FDR=0.0973, Bonferroni-corrected p-value = 0.292) (see Table 1 and Supplementary Table S3). There was no evidence of genome-wide inflation in either analysis (see Figure 2 and Supplementary Table S4). When applying the constraint threshold, there was one top-associated gene that did not meet the 10% BH-FDR threshold and that had somewhat similar results in the schizophrenia cohort (*OLIG2*: CHD uncorrected p-value = 1.39×10^−4^, BH-FDR=0.18; schizophrenia uncorrected p-value = 0.017) (see Supplementary Table S6), indicating questionable validity for CHD. For the genes identified without using the constraint threshold, none had a similar p-value for schizophrenia. Comparing the missense singleton burden in CHD and in gnomAD, genes *NOTCH1*, *BCKDK*, *DHH*, and *KL* displayed similar p-values, but only *NOTCH1* and *BCKDK* passed the BH-FDR 10% threshold (see Supplementary Table S5 and Supplementary Figure S3). These results suggest that for singleton missense gene burden in this study, there was no apparent benefit in restricting to missense-constrained genes.

### Gene-set burden results

Restricting to genes constrained for truncating variants, the gene-set burden analysis (as described in the Materials and Methods) identified one cluster for GO and pathways, and one for MPO, both of which were significant at the sampling FDR < 10%. The FDR approached 1.0 (non-significant) for other clusters (see Table 2). Gene-set sub-clusters were manually identified with the aid of the Cytoscape app EnrichmentMap (13) (see Table 3 and Supplementary Figure S5). The GO and pathway cluster (uncorrected p-value = 5.39×10^−13^, sampling-based FDR = 0) comprised 30 gene-sets, 20 of which were clearly related to VEGF signaling and/or blood vessel development (angiogenesis); *FLT4* was by far the most significant gene (LOF variants N = 7, uncorrected p-value = 9.56×10^−12^), with other genes such as *KDR* (LOF variants N = 2, uncorrected p-value = 0.001), *ZFAND5* (LOF variant N = 1, uncorrected p-value = 0.008) and *WNT5A* (LOF variant N = 1, uncorrected p-value = 0.010) having more modest contributions (see Table 3 and Supplementary Tables S7 and S8). The MPO cluster (uncorrected p-value = 9.64×10^−11^, sampling-based FDR = 0.008) comprised 19 gene-sets, 15 of which corresponded to abnormalities of the cardiovascular system such as abnormal vessel morphology and cardiac-related bleeding in mice (see Table 3 and Supplementary Table S9 and S10). Again, *FLT4* encoding VEGFR3 was the largest contributor, with other genes in the VEGF pathway including *KDR* encoding VEGFR2 and *FOXO1* (LOF N = 1, uncorrected p-value = 0.008) that were previously identified using manual curation (6). The GO pathway and MPO cluster results however also identified other potential candidate genes for TOF associated with functions of *FLT4* that were not identified in the previous study, including *AKAP12*, *PKD1*, *ATF2*, and *EPN1* (Table 3). While other clusters were not significant after multiple test correction, some top-scoring ones had a clear functional or phenotypic relation to CHD (for instance, planar cell polarity in neural tube closure, ranking third for GO and pathways; positive regulation of vascular smooth cell migration, ranking fourth for mouse phenotypes) and included additional promising candidate genes (e.g., *DVL3*, *KIF3A*).

**Table 2).** Top six gene-set clusters for truncating singleton variant burden analyses using Gene Ontology (GO) / pathways and Mouse Phenotype Ontology (MPO), and restricting to constrained genes.

| Gene-set clusters | Observed truncating variants<br>in constrained genes<br>(o/e score in gnomAD <0.35) | p-value | Permutation<br>based FDR |
| --- | --- | --- | --- |
| **GO and pathways** |  |  |  |
| VEGF signaling and blood vessel development | 8 | $5.39 \times 10^{-13}$ | 0 |
| Ion antiporter activity | 5 | 0.0005 | 0.9564 |
| Planar cell polarity pathway involved in neural tube closure | 3 | 0.0013 | 0.9564 |
| Positive regulation of vascular associated smooth muscle cell migration | 4 | 0.0017 | 0.9564 |
| Peptidyl-tyrosine autophosphorylation | 5 | 0.0027 | 0.9564 |
| Protein quality control for misfolded or incompletely synthesized proteins | 3 | 0.0029 | 0.9564 |
| **MPO** |  |  |  |
| Abnormal lymphangiogenesis | 7 | $9.64 \times 10^{-11}$ | 0.008 |
| Abnormal cranial neural crest cell morphology | 3 | 0.0010 | 0.9605 |
| Neuronal cytoplasmic inclusions | 2 | 0.0022 | 0.9605 |
| Absent pharyngeal arches | 4 | 0.0031 | 0.9605 |
| Abnormal CD5-positive T cell number | 2 | 0.0036 | 0.9605 |
| Cochlear ganglion degeneration | 4 | 0.0037 | 0.9605 |

**Table 3).** Gene-set sub-clusters derived from the two gene-set clusters with significant truncating singleton burden from Table 2.

| Most significant composite gene-set sub-clusters | p-value | Genes<sup>1</sup><br>(contributing number of variants,<br>p-value) |
| --- | --- | --- |
| **GO and pathways** |  |  |
| Positive regulation of protein kinase C signaling | $5.39 \times 10^{-13}$ | *FLT4*<sup>2</sup>, *WNT5A* (1, 0.010) |
| Regulation of protein kinase C signaling | $1.17 \times 10^{-12}$ | *FLT4*<sup>2</sup>, *WNT5A* (1, 0.010),<br>*AKAP12* (1, 0.024) |
| VEGF and related pathways, and transmembrane receptor protein kinase activity | $1.68 \times 10^{-12}$ | *FLT4*<sup>2</sup>, *KDR*<sup>3</sup> (2, 0.001) |
| Regulation of blood vessel remodeling, VEGFR3 signaling in lymphatic endothelium, and lung alveolus development | $4.70 \times 10^{-10}$ | *FLT4*<sup>2</sup> |
| Lymph vessel morphogenesis and development | $6.73 \times 10^{-9}$ | *FLT4*<sup>2</sup>, *PKD1* (1, 0.046) |
| Respiratory system process and gaseous exchange | $4.73 \times 10^{-8}$ | *FLT4*<sup>2</sup>, *ZFAND5* (1, 0.008) |
| Endothelial cell proliferation and migration | $6.45 \times 10^{-7}$ | *FLT4*<sup>2</sup>, *KDR*<sup>3</sup> (2, 0.001),<br>*WNT5A* (1, 0.010) |
| **MPO** |  |  |
| Anterior cardinal vein development, abnormal lymph circulation, abnormal lymphatic system physiology, and ascites | $9.64 \times 10^{-11}$ | *FLT4*<sup>2</sup> |
| Abnormal lymphangiogenesis and abnormal lymphatic vessel morphology | $1.55 \times 10^{-10}$ | *FLT4*<sup>2</sup>, *KDR*<sup>3</sup> (2, 0.001) |
| Heart hemorrhage | $4.20 \times 10^{-7}$ | *FLT4*<sup>2</sup>, *KDR*<sup>3</sup> (2, 0.001),<br>*ATF2* (1, 0.036), *PKD1* (1, 0.046) |
| Hemopericardium | $2.33 \times 10^{-6}$ | *FLT4*<sup>2</sup>, *ATF2* (1, 0.036),<br>*PKD1* (1, 0.046) |
| Skin edema and hydrops fetalis | $5.84 \times 10^{-5}$ | *FLT4*<sup>2</sup>, *PKD1* (1, 0.046) |
| Abnormal vitelline vascular remodeling | $2.41 \times 10^{-4}$ | *FLT4*<sup>2</sup>, *KDR*<sup>3</sup> (2, 0.001),<br>*FOXO1*<sup>3</sup> (1, 0.008),<br>*EPN1* (1, 0.015), *TTN* (1, 0.740) |

<sup>1</sup> Genes listed are all those meeting the constraint threshold of o/e less than 0.35, e.g., including genes not reaching significance (e.g., *TTN*).
<sup>2</sup> For each significant gene-set subcluster, for gene *FLT4* the number of variants contributing is 7, and the p-value is $9.56 \times 10^{-12}$ .
<sup>3</sup> Candidate genes previously identified through manual curation methods, relevant to the VEGF pathway, in addition to *FLT4* (6).

Since *FLT4* had such a prominent role in driving the gene-set signal for truncating variants, we repeated the analysis without *FLT4*. No significant gene-set cluster was identified. Similar results were obtained when considering all genes (i.e. without restricting to constrained genes), but the MPO cluster had FDR slightly higher than the 10% threshold (see Supplementary Table S9). For the missense variant analysis, we observed no significant gene-sets, with or without applying the constraint cut-off (see Supplementary Table S11 and S12).

### Detailed *in-silico* analysis of missense variants in *NOTCH1* and other genes

Given that our previous report had focused on truncating variants^6^, we reviewed in detail the singleton missense variants identified, considering amino acid conservation in orthologous vertebrate sequences and *in-silico* predictors (SIFT, PolyPhen2, and Mutation Assessor) (14–16). For *NOTCH1*, this manual review deemed 7 of the 8 ultra-rare missense variants to be either likely deleterious (n=6) or potentially deleterious (n=1). For *BCKDK*, 1 of 4 was likely deleterious, and 1 of 4 potentially deleterious; for *KL*, 3 of 4 were potentially deleterious; for *DHH*, 1 of 3 was likely deleterious and 1 of 3 potentially deleterious; see Supplementary Table S13 for details).

All 8 *NOTCH1* variants identified reside in the extracellular domain of the encoded protein (amino acids 19-1735, see Figure 3), compared to 958 of 1,413 gnomAD v2.1 ultra-rare missense variants (one-sided Fisher’s Exact Test p-value = 0.045, odds ratio = +Inf). Similar to previously reported exome sequencing findings (8), four of these 8 variants alter evolutionarily conserved cysteine residues that establish disulfide bonds, located within the EGF-like repeats or the LNR (Lin12-Notch) domain (17,18). This represents highly significant enrichment compared to such variants from gnomAD v2.1 (23 of 958 variants; one-sided Fisher’s Exact Test p-value = 3.15×10^−5^, odds ratio = 39.8).

**Figure 3).**
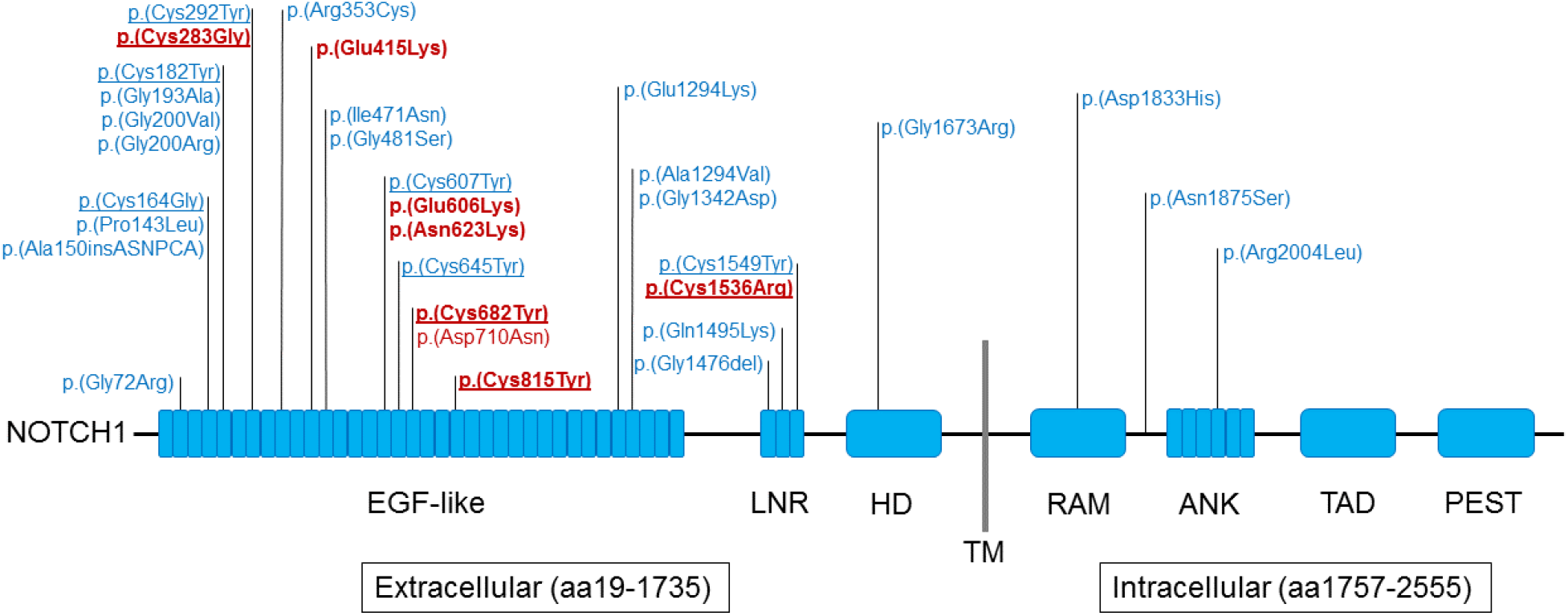
Schematic representation of *NOTCH1* domains (https://www.uniprot.org/uniprot/P46531) and rare variants identified in individuals with tetralogy of Fallot. Findings from the current study involving 8 of 175 probands with TOF are indicated in red font; 24 ultra-rare missense variants from the Page et al. study (8) are indicated in blue font. The seven ultra-rare missense *NOTCH1* variants deemed to be either likely deleterious (n=6) or potentially deleterious (p.(Asn623Lys)) are indicated in bold red font (details in supplementary Table S13). Underline indicates those variants that alter evolutionarily conserved cysteine residues; eight located within the EGF-like repeats domain and two in the LNR (Lin12-Notch) domain. Abbreviations: aa, amino acid; ANK, ankyrin; EGF, epidermal growth factor; HD, heterodimerization domain; LNR, Lin/NOTCH repeats; PEST, sequence rich in proline, glutamic acid, serine, and threonine; RAM, RBP-JK–associated molecule region; TAD, transactivation domain; TM, transmembrane domain (aa1736-1756).

Notably, all 8 of the ultra-rare missense variants in *NOTCH1* were identified within the 175 individuals with TOF, representing 4.6% of those studied. There was significant enrichment for positive family history of CHD compared to the rest of the TOF sample (4 of 8 probands; two-sided Fisher’s Exact Test p-value = 0.003431, odds ratio = 11.49). Details of phenotype and family history are provided for individuals with these 8 *NOTCH1* and 12 other variants in Supplementary Table S14.

## Discussion

In this study, we re-analyzed WGS data available for 231 individuals with CHD, including 175 with TOF, to extend previously published results^6^ using a statistical method modified to suit such case-only data. By rescaling *de novo* mutation probabilities for singleton variants, we adapted a burden test originally developed for *de novo* variants, and tested truncating and missense singleton variants separately for increased burden in genes and in functionally relevant gene-sets.

Previous results suggested that ultra-rare nonsynonymous variants make an important contribution to the genetic etiology of CHD, especially to TOF (5,6,8). Since constrained genes may be more likely to contribute to disease, in order to maximize power, we performed multiple test correction for all genes, and only for genes passing a constraint threshold. We assessed the validity of our results by ensuring the absence of inflation when considering the burden test p-value distribution. In addition, we compared burden results for CHD to a schizophrenia WGS data-set processed in the same way (including variant calling and QC), to help identify potential artifacts. Finally, we also retested burden by comparing results to gnomAD singletons that had been processed in the same way with respect to variant effect and singleton definition.

### Gene burden

Two genes passed a very stringent significance threshold of 0.01 after Bonferroni correction: *FLT4* for truncating variants and *NOTCH1* for missense variants. Burden significance for these genes was highly specific to CHD, compared to an unrelated schizophrenia sample, was confirmed by the gnomAD singleton comparison analysis, and involved only individuals with TOF. The results are consistent with exome sequencing results from an independent study of 829 individuals with TOF, analyzed using a different approach, where excess of ultra-rare deleterious variants was reported to be genome-wide significant (p-value ≤ 5×10^−8^) for these two genes (8) and for *FLT4* in another study restricted to LOF variants (5). These exome sequencing results serve to both help validate our burden test methodology and provide independent replication, further cementing these genetic findings for TOF. Collectively, the findings support study designs that focus on TOF.

For truncating variants, restricting to constrained genes did not result in identifying any other significant genes, even when considering a more inclusive significance threshold of BH-FDR < 10%. Including all genes resulted in one other gene that passed BH-FDR < 10%, *CLDN9* (Claudin 9). *CLDN9* burden was not however confirmed by the comparison to gnomAD singleton variants and the gene lacks evidence for involvement in cardiovascular development, thus at present we consider this result to be likely artifactual. The results suggest that, in order to limit such artifacts, considering only LOF-constrained genes may be especially important when a well matched data-set (here, schizophrenia WGS) is not available. For example, artifacts can arise if *de novo* mutation probabilities are derived from WGS data that were processed differently than the data available for the case-only cohort (e.g., different variant calling pipeline, QC filters, and principal transcripts). Also, *denovolyzer* probabilities were generated for exome analysis and adjusted for sequencing depth, thus artifacts may arise in WGS studies where sequencing depth is greater. In contrast, for missense variants, testing only genes passing a missense constraint threshold did not appear to be beneficial. This is perhaps because missense constraint tends to be a characteristic of specific protein regions rather than the full gene product and this is not adequately modelled by gnomAD constraint indexes.

For ultra-rare missense variants, we identified slightly different sets of significant genes (BH-FDR < 10%) when considering only constrained genes or all genes. *BCKDK* (Branched-chain keto acid dehydrogenase kinase) was identified in both analyses; *DHH* (Desert hedgehog signaling molecule) was identified only in the constrained analysis, and *KL* (Klotho) only in the all-gene analysis. All displayed a similar p-value in the gnomAD singleton comparison, and only *BCKDK* passed BH-FDR < 10%. *BCKDK* is a negative regulator of the branched-chain amino acids catabolic pathways. In mice, complete *BCKDK* loss of function causes reduced size and neurological abnormalities (19). In humans, recessive *BCKDK* loss of function causes recessive ‘Branched-chain ketoacid dehydrogenase kinase deficiency’ (OMIM: 614923), characterized by autism, epilepsy, intellectual disability, and risk of developing schizophrenia (20). Alterations of branched-chain amino acid metabolism have been described in relation to heart failure (21), however, there is no evident link between *BCKDK* and CHD. *DHH* is required for Sertoli cells development and peripheral nerve development: male Dhh-null mice are sterile and fail to produce mature spermatozoa; in addition, peripheral nerves are highly abnormal (22,23). These features are mirrored by the human recessive disorder ‘46XY partial gonadal dysgenesis, with minifascicular neuropathy’ (OMIM: 607080), whereas the recessive disorder ‘46XY sex reversal 7’ (OMIM: 233420) does not present peripheral nerve abnormalities (24,25). No cardiac abnormalities were reported for these disorders. However, *DHH* was also proposed to contribute to promoting ischemia-induced angiogenesis by ensuring peripheral nerve survival (26). In humans, *KL* was previously proposed as a candidate gene for TOF because of one patient with a broader 13q13 deletion and another patient with a narrower deletion at the same locus disrupting *KL* and *STARD13* (27,28). Deficiency of *Kl* in mice has profound systemic effects, resulting in a phenotype reminiscent of human ageing and characterized by reduced lifespan, stunted growth, skeletal abnormalities, vascular calcification and atherosclerosis, cognitive impairment and other organ alterations (29). Conversely, over-expression of *Kl* in mice protects against cardiovascular disease. In addition, *Kl* is involved in the regulation of several pathways, including VEGF and Wnt (30). However, considering the evidence reviewed above and that these three genes present a lower fraction of singleton predicted deleterious missense variants compared to *NOTCH1*, caution is needed when considering these genes as candidates for CHD/TOF. Replication in larger cohorts and/or experimental follow-up is required.

### Functional gene-sets and candidate genes

Since constrained genes would be expected to have a low rate of ultra-rare variation and thus present power challenges in a cohort of this size, in addition to assessing variants by gene, we also pooled variants by functional gene-sets and mouse ortholog phenotypes. To correct for strong correlations introduced by highly overlapping gene-sets, which may result in inflated significance after multiple test correction, we used a greedy clustering procedure to group highly overlapping gene-sets; we then performed multiple test correction using a sampling-based FDR. Reassuringly, given our previously published results (6), the gene-set burden analysis for truncating singleton variants yielded a cluster corresponding to the VEGF pathway and blood vessel development (FDR = 0), and also a cluster corresponding to abnormal vasculature (FDR = 0.008). As expected (6), *FLT4* was the main gene driving these results. We additionally identified other genes that were only nominally significant, but had suggestive functional or phenotypic evidence and could achieve genome-wide significance in a larger cohort.

Although we had previously identified some of these genes (*KDR* and *FOXO1*) (6), *WNT5A* (Wnt family member 5A) and *ZFAND5* (zinc finger AN1-type containing 5), were identified only in this re-analysis and appear particularly promising candidates for TOF/CHD. *ZFAND5* is transcriptionally activated by the platelet-derived growth factor (PDGF) pathway (31), and is reported to be a member of the FoxO family signaling pathway by the NCI-Nature PID pathway database. Mice homozygous for a *Zfand5* null mutation die postnatally due to widespread bleeding, caused by loss of vascular smooth muscle cells (31). Mice homozygous for a *Wnt5a* null allele die perinatally, with reduced growth and multiple organ system developmental abnormalities; notably, the heart presents outflow tract defects and *Wnt5a* loss disrupts second heart field cell deployment; heterozygous mice are apparently normal (32–34). Functional experiments in mice showed that *Wnt5a* contributes to the vascular specification of cardiac progenitor cells and a role in pressure overload-induced cardiac dysfuncion (35,36). In humans, heterozygous missense or homozygous loss of function variants in *WNT5A* are associated with ‘Robinow syndrome’ (OMIM: 180700) (37), which is characterized by short stature, macrocephaly, delayed bone age and limb shortening, reproductive system and kidney abnormalities (38). CHD and specifically right ventricular outlet obstruction are present in a fraction of the cases (39). The results also suggested other constrained genes not previously identified, with evidence that supports a role in the VEGF pathway or other complementary mechanisms for TOF, from human (e.g., *AKAP12*), mouse (e.g., *EPN1*, *ATF2*) or both (*PKD1*) (Table 3) derived gene-sets (40–44).

We note that, collectively, genes *NOTCH1*, *FLT4*, *ZFAND5* and *WNT5A* present singleton variants in probands with TOF but no other-CHD, representing significant enrichment (Fisher test two-sided p-value = 0.01494) compared to background total sample. These account in total for about 11% of the individuals with TOF studied (see Supplementary Table 14).

One may wonder why certain VEGF pathway genes that were previously implicated in TOF using manual curation of this data-set (6) were not found in the gene-set analysis in the current study. There are several possible reasons. *BCAR1* was implicated by structural variation (thus not analyzed in the current study), *VEGFA* does not have a defined *de novo* mutation probability in *denovolyzer*, *FGD5* and *PRDM1* are not associated to any VEGF-related gene-sets among the GO and pathway gene-sets used for this analysis, and *IQGAP1* was present only in a VEGF-related gene-set that did not contain *FLT4* and thus did not achieve significance. If these genes were also included, this would account for around 14% TOF individuals.

### Advantages and limitations

Results from several published studies suggest that analyzing ultra-rare variant burden is a suitable strategy for CHD, and especially for TOF (5,6,8), given a genetic architecture characterized by rare variants of large effect, reduced penetrance and oligogenic contributions. The method we adopted enables testing of ultra-rare genetic variant burden in a case-only cohort, without having access either to parents to determine variant *de novo* status or to matched controls for case-control analysis. This would be a relatively common circumstance for many studies, especially of rare and under-funded conditions.

In our study design, we attempted to address issues that can produce artifacts, such as mismatch of the variant calling, and/or processing pipelines, between those used for the disease data-set and for the data-set supporting the calculation of *de novo* probabilities. We had the advantage of access to a similarly sized sequencing data-set for an unrelated disease, processed in the same way, to aid in identifying potential artifacts that may not be available for future applications of this statistical burden method. Although we observed that restricting the burden analysis to genes constrained for truncating variants may help minimize such artifacts, there was no apparent advantage using constraint for missense variants, and we note that the findings may be disease or study specific.

Our primary gene burden analysis was based on *de novo* mutation probabilities rescaled to match incidence of singleton variants, with probabilities defined separately for truncating and missense variants. As a further confirmatory analysis to the unrelated cohort with similar WGS data, we compared the singleton burden in CHD to that in gnomAD. Additional analyses using a benchmark are required to establish whether one of these two methods is superior, in terms of power and minimizing artifacts. The advantage of using gnomAD singletons is that, while variant calling pipelines cannot be matched, other downstream processes like annotation can be matched to the disease data-set of interest.

All results were limited by the size of the cohort available with WGS data. There were also limitations to the design that are applicable to all analyses using gene-sets, including the lag in updating bioinformatics databases (45) such as GO and MPO. These limitations could have had an impact on that fact that there was no significant gene-set identified for missense variants. Also, although the method identified highly relevant gene-set clusters for singleton truncating variants, *FLT4* played a disproportionately large role in the analysis, likely influencing the fact that the relatively few novel candidate genes identified largely converged on the VEGF pathway. For other disorders that are even more genetically heterogeneous, the results suggest that optimizing the analysis method at the gene-set level may be essential in order to identify significant results (45,46). As for all studies using statistical methods to identify potential disease candidate genes, additional experimental work would be required to conclusively implicate genes.

Sample size limitations, and the genetic architecture of TOF, also likely influenced the gene-based analysis findings. Nonetheless, two genes passed a stringent Bonferroni correction, *FLT4* (implicated by truncating variants) and *NOTCH1* (implicated by missense variants), consistent with previous findings reported in two (5,8) and one (8) independent exome sequencing studies, respectively. Notably, results of the current study indicated higher yields of ultra-rare variants in these genes, perhaps related to differences in design and methods, including sequencing (WGS) expected to have more uniform and complete coverage of coding regions, and perhaps the use of an adult sample that would enrich for variants associated with survival. Future meta-analyses using this and other data-sets, or studies using genetically relevant subsets of patients to reduce heterogeneity, could reveal additional candidate genes with rare variants for TOF. In particular, several gene-set clusters did not pass the multiple test correction yet appeared highly promising, and could achieve significance in an expanded cohort.

Non-coding variants and structural variants, which can be detected using whole genome data (47), were not studied. For rare non-coding variants, access to large samples of whole genome data may offer interesting opportunities, especially if analyzing better understood functional elements like promoters. The lack of a published mutation probability model for promoters could be circumvented by performing only the gnomAD comparison analysis. For structural variants, although reliably detectable using WGS (in contrast to exome sequencing), lack of a *de novo* mutation model, and greater variability in variant calling pipelines, would prevent a direct gnomAD comparison and represent major barriers at present.

## Conclusions

The gene burden analysis method used, including a stringent Bonferroni correction, confirmed that genes *FLT4* with ultra-rare truncating variants, and *NOTCH1* with ultra-rare deleterious missense variants, are implicated in the etiology of TOF. The significant enrichment of *NOTCH1* missense variants in the extracellular domain, and specifically altering cysteine residues forming disulfide bonds, was also confirmed. Despite the small sample size, gene-set analysis identified ultra-rare truncating variants in novel candidate genes, including *ZFAND5* and *WNT5A*, as potentially implicated in the etiology of TOF. Other novel genes identified provide further confidence in the importance of the VEGF pathway to TOF. While several of these candidate genes are compelling, with supportive data from known functions and animal model phenotype, additional experimental work and/or replication in other data-sets are required to appreciate their potential role in the etiology and pathogenesis of TOF.

## Materials and Methods

### Study participants and genome sequencing

This study was authorized by the Research Ethics Boards at the University Health Network (REB 98-E156) (http://www.uhn.ca), and Centre for Addiction and Mental Health (REB 154/2002) (http://www.camh.ca). Written consent was obtained from all participants or their legal guardians.

We performed genome sequencing on 231 probands (175 TOF, 49 transpositions of the great arteries, 7 other CHD). DNA was sequenced on the Illumina HiSeq X system (https://www.illumina.com/systems/sequencing-platforms/hiseq-x.html) at The Centre for Applied Genomics (TCAG) (http://www.tcag.ca). Libraries were amplified by PCR prior to sequencing. Libraries were assessed using Bioanalyzer DNA High Sensitivity chips and quantified by quantitative PCR using Kapa Library Quantification Illumina/ABI Prism Kit protocol (KAPA Biosystems). Validated libraries were pooled in equimolar quantities and paired-end sequenced on an Illumina HiSeq X platform following Illumina’s recommended protocol to generate paired-end reads of 150 bases in length.

### Variant Calling, Annotation, and Truncating and Missense Variant extraction

#### Variant Calling

The paired FASTQ reads were mapped to the GRCh37 reference sequence using the BWA-backtrack algorithm (v0.7.12), and SNV and small indel variants were called using GATK (v3.7) according to GATK Best Practices recommendations (48,49).

#### Variant Annotation

Variant calls were annotated using a custom pipeline based on ANNOVAR (July 2017 version) (50). Allele frequencies were derived from 1000 genomes (Aug. 2015 version) (51), ExAC (Nov. 2015 version) (7), and gnomAD (Mar. 2017 version) (52).

#### Classification of variants by truncating and missense effect

Truncating variants (labelled as *LOF* for *loss of function*) comprised frameshift insertions/deletions, alterations of the highly conserved intronic dinucleotide at splice sites and substitutions creating a premature stop codon (stop gain). Missense variants are substitutions of amino acids.

### Variant filters based on quality, allele frequency and effect

#### Allele frequency and singleton filter

The burden test adopted in this study was originally developed for *de novo* variants, but we argue that ultra-rare variants are not present in the general population and are likely to have arisen recently from *de novo* mutations transmitted to the progeny. We defined ultra-rare singleton variants as appearing only once in the CHD WGS data-set and never in population reference data-sets (1000 genomes, ExAC, and gnomAD).

#### Low quality filter

We removed variants deemed to be low quality, which met at least one of these criteria: (i) low sequencing depth (DP ≤ 10); (ii) low alternate allele read fraction or low genotype quality (for heterozygous variants, alt_fraction < 0.3 or GQ ≤ 99, for homozygous variants, alt_fraction < 0.8 or GQ ≤ 25).

#### Frameshift indel filter

For each subject, whenever we found multiple indels on the same gene, we removed them from the variants list if their cumulative size was a multiple of 3. Otherwise, we kept one of the indels as a representative and removed the rest.

#### Splice site alteration filter

For insertions overlapping splice sites, we considered them as truncating variants only if the alternate allele sequence did not encode a canonical AG/GT intronic dinucleotide.

#### Principal transcript effect filter

We used the APPRIS database (assembly version: GRCh37, gene dataset: RefSeq105, Oct. 2018) to identify principal transcript isoforms (53) and retained only variants with an effect on a principal transcript. APPRIS principal transcript identification is based on conservation, presence of protein domains and other coding sequence characteristics.

#### Final singleton counts

We considered maximum only one singleton missense or truncating variant per gene per subject, such that, for each variant type, the count of singleton variants in a given gene equals the count of subjects with at least one singleton variant in that given gene.

### Gene burden analysis

#### De novo mutation probabilities

We obtained *de novo* mutation probabilities for each gene from denovolyzeR (http://denovo-lyzer.org/) (54). 1000 Genomes intergenic regions that are orthologous between humans and chimps were used to derive mutation probabilities. The probabilities were based on substitution type, trinucleotide context and other genome structure characteristics; in addition, they were adjusted for exome sequencing depth (55).

#### Rescale de novo mutation probability for singleton variants

Since the original mutation probabilities were estimated for *de novo* variants, we applied a multiplicative global scaling factor (SF), defined in equation [1], to obtain new rescaled probabilities *P_exp,(LOF or Missense),g_*; the scaling factor *SF* is computed so that the number of predicted and observed singleton variants match.

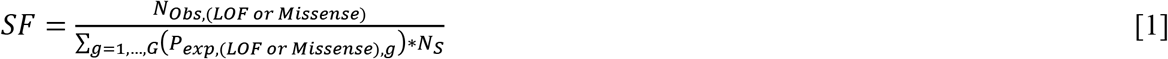

where *N_Obs,(LOF or Missense),g_* is the number of all observed truncating or missense singleton variants; the denominator corresponds to the number of expected singletons using the original unscaled probabilities: *G* is the total number of genes for which there is a defined mutation probability (and optionally, that pass gnomAD constraint cut-offs); *P_exp,(LOF or Missense),g_* is the expected *de novo* mutation probability for gene g with respect to truncating or missense variants; and *N_S_* is the number of subjects in the study.

#### Binomial test

Singleton truncating and missense burden was tested using a one-sided binomial test comparing observed to expected singleton rates, where expected singleton rates correspond to the rescaled mutation probabilities. The alternative hypothesis is defined as P_success_>N_success_/N_trials_, i.e. that the observed singleton rate for a given gene exceeds the expected rate based on rescaled mutation probabilities.

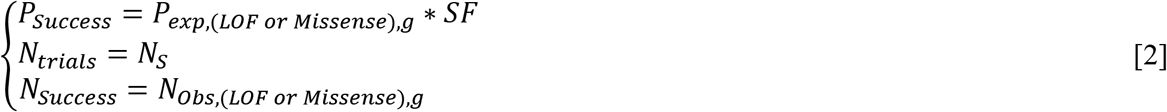

where *N_Obs,(LOF or Missense),g_* denotes the number of observed truncating or missense singleton variants for gene g. Note that, for simplicity, we used singleton variant counts in equation [2], but since we considered maximum only one singleton truncating or missense variant per subject per gene, the singleton truncating or missense variant count per gene is equivalent to the count of subjects with at least one truncating or missense singleton variant in that gene.

#### gnomAD comparison analysis

SNVs and indels data were obtained from the gnomAD v2.1.1 database, comprising WES (125,748 subjects) and WGS (15,708 subjects), after restricting to the interval list (hg19-v0-wgs_evaluation_regions.v1.interval_list) used to generate the Exome Calling Intervals VCF file (gnomad.genomes.r2.1.1.exome_calling_intervals.sites.vcf.bgz). Singleton variants were identified by using the allele counts provided in the gnomAD VCF file. Singletons were annotated using the same ANNOVAR-based pipeline, followed by the same effect filters as in the main analysis (including the selection of the same principal transcript) and finally categorized as truncating or missense. Genes were tested for singleton burden by comparing CHD WGS singletons to gnomAD singletons using a two-sided Fisher’s Exact Test, and specifically by constructing the 2×2 contingency matrix with counts: (a) CHD singletons in the gene of interest, (b) CHD singletons in other genes, (c) gnomAD singletons in the gene of interest, (d) gnomAD singletons in other genes; truncating and missense singletons were tested separately. For CHD, only maximum one singleton per subject was considered (as in the main analysis).

#### Multiple test correction

For gene burden analyses, multiple test correction was performed using the Benjamini-Hochberg False Discovery Rate (*BH-FDR*), as implemented in the R function *p.adjust*, and Bonferroni correction, by multiplying the p-value by the number of genes tested. For both corrections, we considered all genes with a defined probability, or all genes with a defined probability and passing constraint cut-offs (o/e gnomAD score < 0.35 for truncating variants and o/e gnomAD score < 0.75 for missense variants).

### Gene-set burden analysis

#### Gene-set resources

Gene-sets were derived from Gene Ontology (GO) annotations as provided by the Bioconductor package org.Hs.eg.db v3.5 (9), BioCarta pathways (http://cgap.nci.nih.gov/Pathways/BioCarta-Pathways/), KEGG pathways (http://www.genome.jp/kegg/) retrieved using the KEGG API (10), REACTOME pathways (11), and National Cancer Institute (NCI) pathways (https://cactus.nci.nih.gov/download/nci/). Gene-sets corresponding to phenotypes of mouse orthologues were derived from MPO gene annotations as provided by MGI (12).

#### Gene-set filters

We retained only the gene-sets with more than 5 genes and less than 100. Smaller gene-sets are detrimental for power. Larger gene-sets are usually removed because they are overly general. Considering the specific gene-level burden signal distribution observed for this data-set, characterized by the presence of two “highly concentrated” burden genes (*FLT4* and *NOTCH1*), some larger gene-sets could exceed the expected singleton rate just because of the presence of one of these two genes. In addition, larger gene-sets are less suitable for the binomial test strategy, since they are more likely to present with more than one singleton variant per subject and to contain genes with heterogeneous mutation probabilities, which is detrimental when pooling counts (13).

For the analyses using a given gene constraint cut-off, we removed gene-sets with less than two genes passing the constraint cut-offs.

#### Binomial Test

For the gene-set analysis, we used a binomial test (equation [3]) to compare the number of observed and expected singleton variant in the gene-set, similar to the gene burden analysis. We additionally ensured not to count more than one truncating or missense singleton per gene-set per subject.

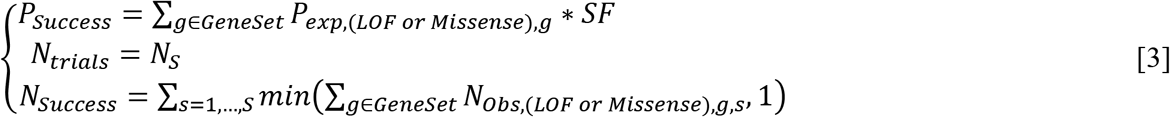

where *GeneSet* represents the set of all genes in a particular gene-set; S are the study subjects; and *N_Obs,(LOF or Missense),g,s_* is the number of observed missense or truncating singleton variants in a particular gene for subject S.

#### Greedy step-down aggregation method to correct for gene-set correlations

We addressed the problem of gene-set correlations, which are introduced by large gene overlaps between related gene-sets, by using a greedy step-down clustering approach, similar to what was adopted for highly correlated CNV locus gene testing in the *Marshall et al.* study (46). The algorithm follows these steps, starting from an input list of gene-sets sorted by the singleton burden binomial p-value:

1. Select the gene-set with the most significant p-value (i.e. the smallest p-value);
2. Identify other gene-sets that are highly correlated to the selected gene-set, using the *Jaccard* similarity:

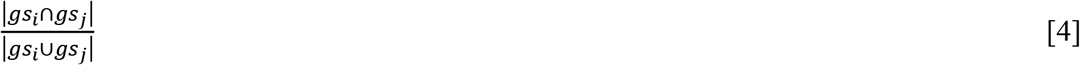

where *gs_i_* and *gs_j_* are the sets of singleton variants for gene-sets i and j, respectively. | | is the number of singleton variants in the corresponding set.
3. Cluster gene-sets that have *Jaccard* similarity > 0.5 with the selected gene-set; these gene-sets will not be considered for the multiple test correction calculation, only the selected gene-set will be used (i.e. the p-value from the selected gene-set will be used as the p-value for the gene-set cluster). Finally, remove the selected gene-set and its clustered gene-sets from the sorted list.

Steps 1-3 are executed until no gene-set is left in the list.

### Resampling-based FDR

Observed missense or truncating singleton variants are resampled based on each gene’s rescaled mutation probability (equation [1]), while maintaining the same total number of observed missense or truncating singleton variants. After this step, gene-sets are tested as described in the previous section. Finally, for each given p-value threshold *p*, the FDR is calculated as follows, considering only gene-sets selected by the greedy step-down aggregation procedure:

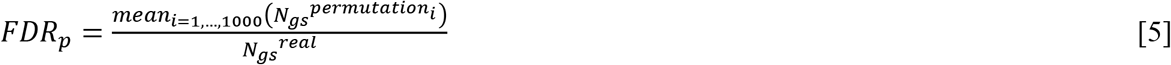

where *FDR_p_* is the FDR q-value for a given p-value threshold *p*, *N_gs_^real^* is the number of gene-sets with binomial p-value ≤ p, and 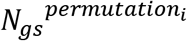 corresponds to the number of gene-sets with binomial p-value ≤ *p* at iteration *i*. As stated in the formula, we used 1,000 sampling iterations.

## Supporting information

Supplementary Figure S1) Relation between gnomAD genetic constraint indexes.

Supplementary Figure S2) Relation between the number of singleton truncating variants per gene in the CHD data-set and in gnomAD

Supplementary Figure S3) Relation between the number of singleton missense variants per gene in the CHD data-set and in gnomAD

Supplementary Figure S4) QQ-plots and p-value CHD/SZ scatterplot for the gene burden analysis restricted to constrained genes

Supplementary Figure S5) Cytoscape enrichment map for the gene-sets with significant burden of singleton truncating variants in constrained genes

Supplementary Table S1) Ultra-rare singleton variants observed for 231 CHD samples including gnomAD o/e constraint scores

Supplementary Table S2) Gene burden statistics for CHD singleton truncating variants

Supplementary Table S3) Gene burden statistics for CHD singleton missense variants

Supplementary Table S4) Truncating variants observed in both SZ and CHD cohorts

Supplementary Table S6) Missense Variants observed in both SZ and CHD cohorts

Supplementary Table S7) Burden statistics for GO and pathway gene-set clusters and truncating variants in constrained genes

Supplementary Table S8) Burden statistics for GO and pathway gene-sets and singleton truncating variants in constrained genes

Supplementary Table S9) Burden statistics for mouse phenotype gene-set clusters and singleton truncating variants in constrained genes

Supplementary Table S10) Burden statistics for mouse phenotype gene-sets and singleton truncating variants in constrained genes

Supplementary Table S11) Burden statistics for GO and pathway gene-set clusters and singleton missense variants in constrained genes

Supplementary Table S12) Burden statistics for GO and pathway gene-sets and singleton missense variants in constrained genes

Supplementary Table S13) NOTCH1, BCKDK, DHH and KL missense variants details

Supplementary Table S14) Patient phenotype and family history for selected deleterious missense and truncating variants

## Acknowledgments

We thank the patients and their families for participating in this study.

## Legends to Supplementary Figures and Tables

### Supplementary Figures

**Supplementary Figure 1**

**Relation between gnomAD genetic constraint indexes.**

(A) Relationship between pLI (x axis, discretized in three bins) and the ratio of observed/expected (o/e) truncating variants (y axis). pLI > 0.9 has often been used as haploinsufficiency cutoff for clinical variant interpretation, and gnomAD suggests using the upper bound of the o/e confidence interval < 0.35 for a similar use. We preferred using a point estimate <= 0.35 to be more inclusive, i.e. including genes with more moderate haploinsufficiency. For our analysis, we have considered genes with o/e score < 0.35. (B) Relationship between the missense constraint z-score (x axis, discretized in two bins) and the ratio of observed/expected missense variants (y axis). For our analysis, we have considered genes with o/e score < 0.75, which roughly corresponds to a z-score > 2, which in turn corresponds to a constraint p-value of 0.02275.

**Supplementary Figure 2**

**Relation between the number of singleton truncating variants per gene in the CHD data-set and in gnomAD.**

The distribution (across genes) of the number of singleton truncating variants per gene is shown as an overlaid boxplot and violin plot for singletons in gnomAD (x axis), stratified by the the number of singleton variant in the CHD data-set (y axis); each dot represents a gene. The dashed line represents the linear regression predictions, which are appear unreliable because of outliers and the small number of unique CHD singleton counts. Only *FLT4* has 7 truncating singletons, but the trend for other strata suggests that this is in large excess of singletons observed in gnomAD. Note that CHD singletons are not observed in gnomAD, whereas gnomAD singletons are observed only once in gnomAD.

**Supplementary Figure 3**

**Relation between the number of singleton missense variants per gene in the CHD data-set and in gnomAD.**

The distribution (across genes) of the number of singleton missense variants per gene is shown as an overlaid boxplot and violin plot for singletons in gnomAD (x axis), stratified by the the number of singleton variant in the CHD data-set (y axis); each dot represents a gene. The dashed line represents the linear regression predictions, which appear robust. *KL* and *DHH* overlap with the lowest percentiles of the distribution, whereas *BCKDK* is lower than any observed value; only NOTCH1 has 8 missense singletons, but the trend for other strata suggests that this is in excess of singletons observed in gnomAD. Note that CHD singletons are not observed in gnomAD, whereas gnomAD singletons are observed only once in gnomAD.

**Supplementary Figure 4**

**QQ-plots and p-value CHD/SZ scatterplot for the gene burden analysis restricted to constrained genes.**

(A) and (C) show the quantile-quantile (QQ) plots for gene burden p-values obtained for truncating singletons restricted to constrained genes (gnomAD o/e < 0.35, A) or missense singletons restricted to constrained genes (gnomAD o/e < 0.75, B). Only a few genes present p-values deviating from the null distribution, suggesting absence of systematic p-value inflation. (B) and (D) show scatterplots of the nominal p-values obtained for the gene burden analysis of truncating or missense singleton variants in constrained genes, comparing CHD (y axis) versus schizophrenia (x axis). The most significant genes for CHD are typically not significant for SZ, suggesting the absence of systematic confounders.

**Supplementary Figure 5**

**Cytoscape enrichment map for the gene-sets with significant burden of singleton truncating variants in constrained genes.**

An enrichment map visualizes gene-sets as a network based on their overlaps. Nodes correspond to gene-sets from the gene-set cluster with significant (FDR < 10%) burden for truncating singleton variants in constrained genes, and edges correspond to the degree of overlap between gene-sets. Nodes are colored based on the burden nominal p-value, with darker red corresponding to more significant gene-sets. Edge thickness is proportional to the jaccard index obtained by considering singleton truncating variants as set elements; only edges corresponding to jaccard index > 0.5 are displayed. Gene-set sub-clusters are suggested by automated network layout. Gene Ontology and pathways (A) are shown separately from mouse phenotypes (B).

### Supplementary Tables

**Supplementary Table 1**

**Ultra-rare singleton variants observed for 231 CHD samples including gnomAD o/e constraint scores**

**Supplementary Table 2**

**Gene burden statistics for CHD singleton truncating variants**

**Supplementary Table 3**

**Gene burden statistics for CHD singleton missense variants**

**Supplementary Table 4**

**Truncating variants observed in both SZ and CHD cohorts**

**Supplementary Table 5**

**Gene burden statistics obtained by comparing CHD singletons to gnomAD singletons and tested using Fisher’s Exact Test**

**Supplementary Table 6**

**Missense Variants observed in both SZ and CHD cohorts**

**Supplementary Table 7**

**Burden statistics for GO and pathway gene-set clusters and truncating variants in constrained genes**

**Supplementary Table 8**

**Burden statistics for GO and pathway gene-sets and singleton truncating variants in constrained genes**

**Supplementary Table 9**

**Burden statistics for mouse phenotype gene-set clusters and singleton truncating variants in constrained genes**

**Supplementary Table 10**

**Burden statistics for mouse phenotype gene-sets and singleton truncating variants in constrained genes**

**Supplementary Table 11**

**Burden statistics for GO and pathway gene-set clusters and singleton missense variants in constrained genes**

**Supplementary Table 12**

**Burden statistics for GO and pathway gene-sets and singleton missense variants in constrained genes**

**Supplementary Table 13**

**NOTCH1, BCKDK, DHH and KL missense variants details**

**Supplementary Table 14**

**Patient phenotype and family history for selected deleterious missense and truncating variants**

