## Supplementary figures and images for "Genes and pathways implicated in tetralogy of Fallot revealed by ultra-rare variant burden analysis in 231 genome sequences"

### Supplementary Figure S1) Relation between gnomAD genetic constraint indexes.

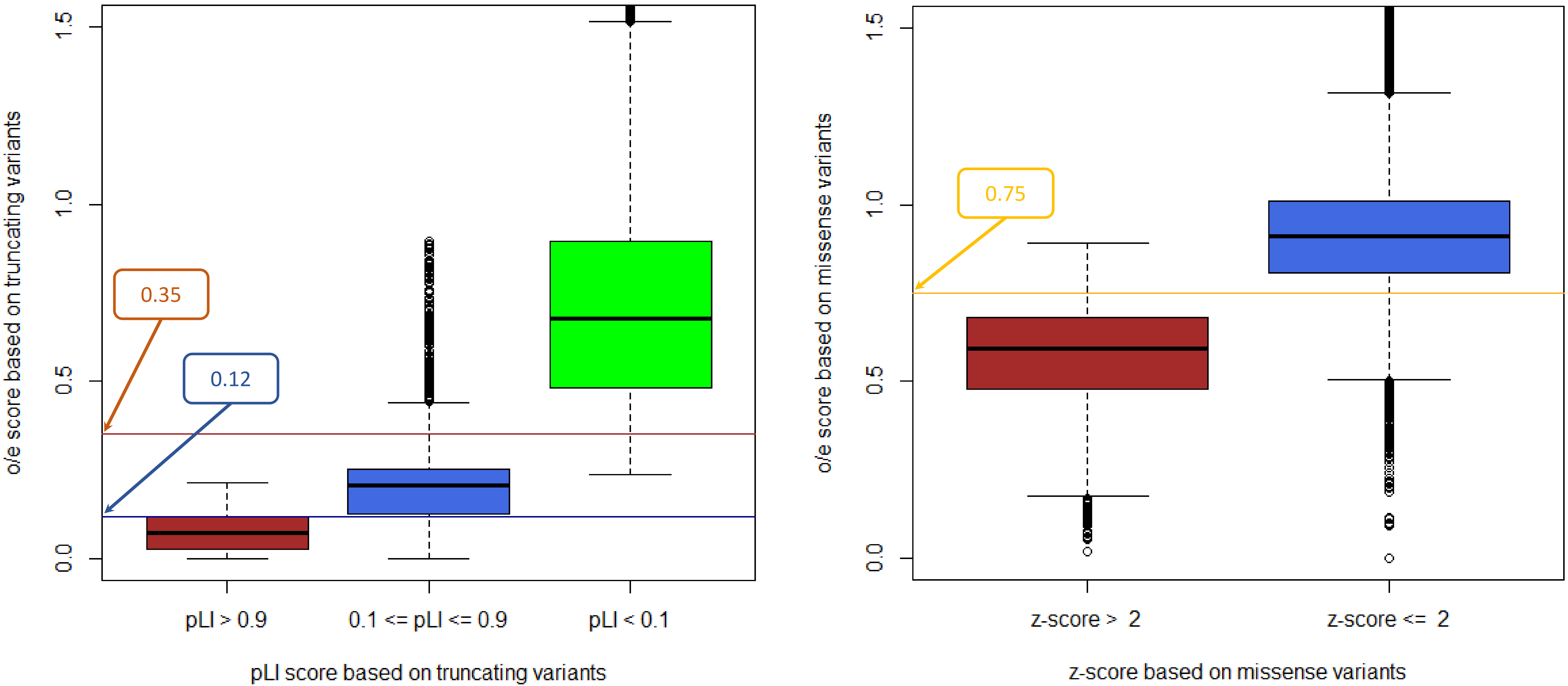

### Supplementary Figure S2) Relation between the number of singleton truncating variants per gene in the CHD data-set and in gnomAD

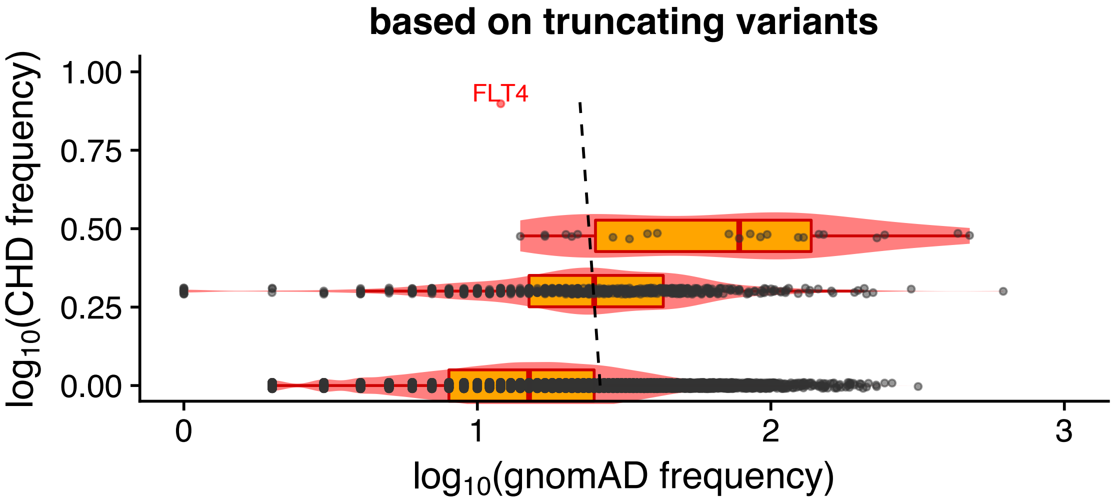

### Supplementary Figure S3) Relation between the number of singleton missense variants per gene in the CHD data-set and in gnomAD

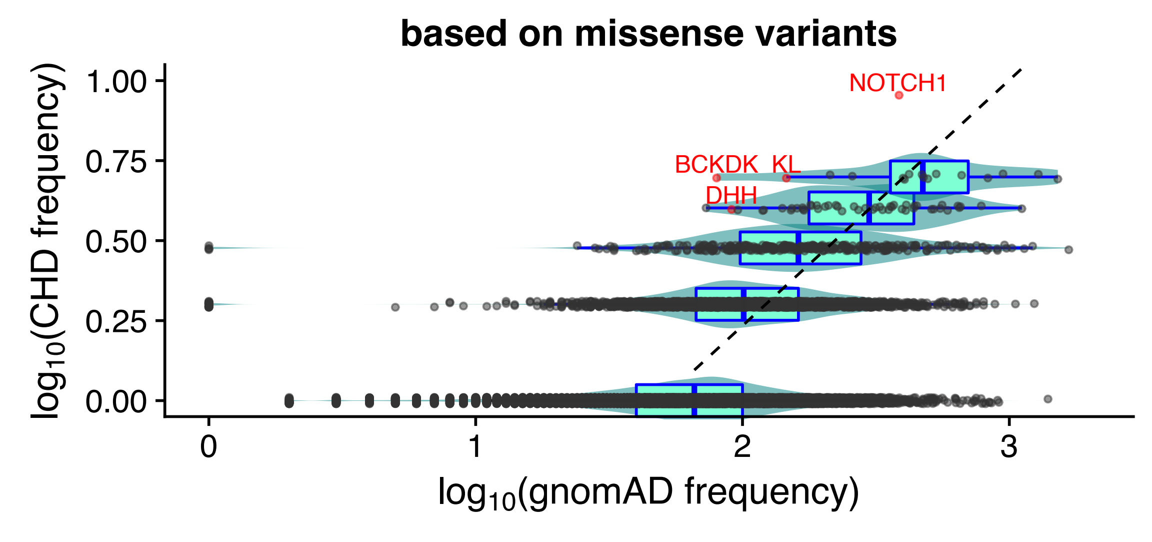

### Supplementary Figure S4) QQ-plots and p-value CHD/SZ scatterplot for the gene burden analysis restricted to constrained genes

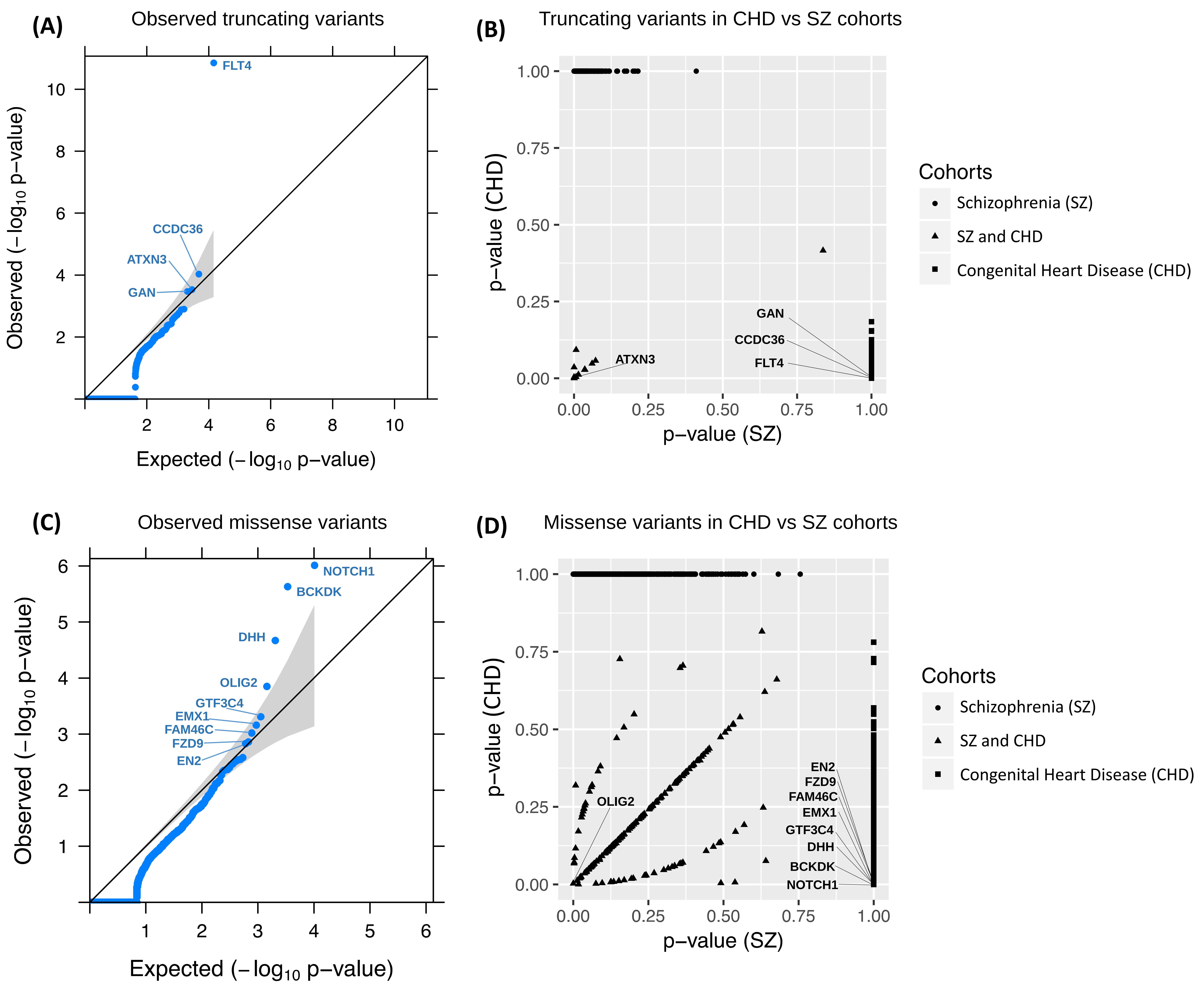

### Supplementary Figure S5) Cytoscape enrichment map for the gene-sets with significant burden of singleton truncating variants in constrained genes

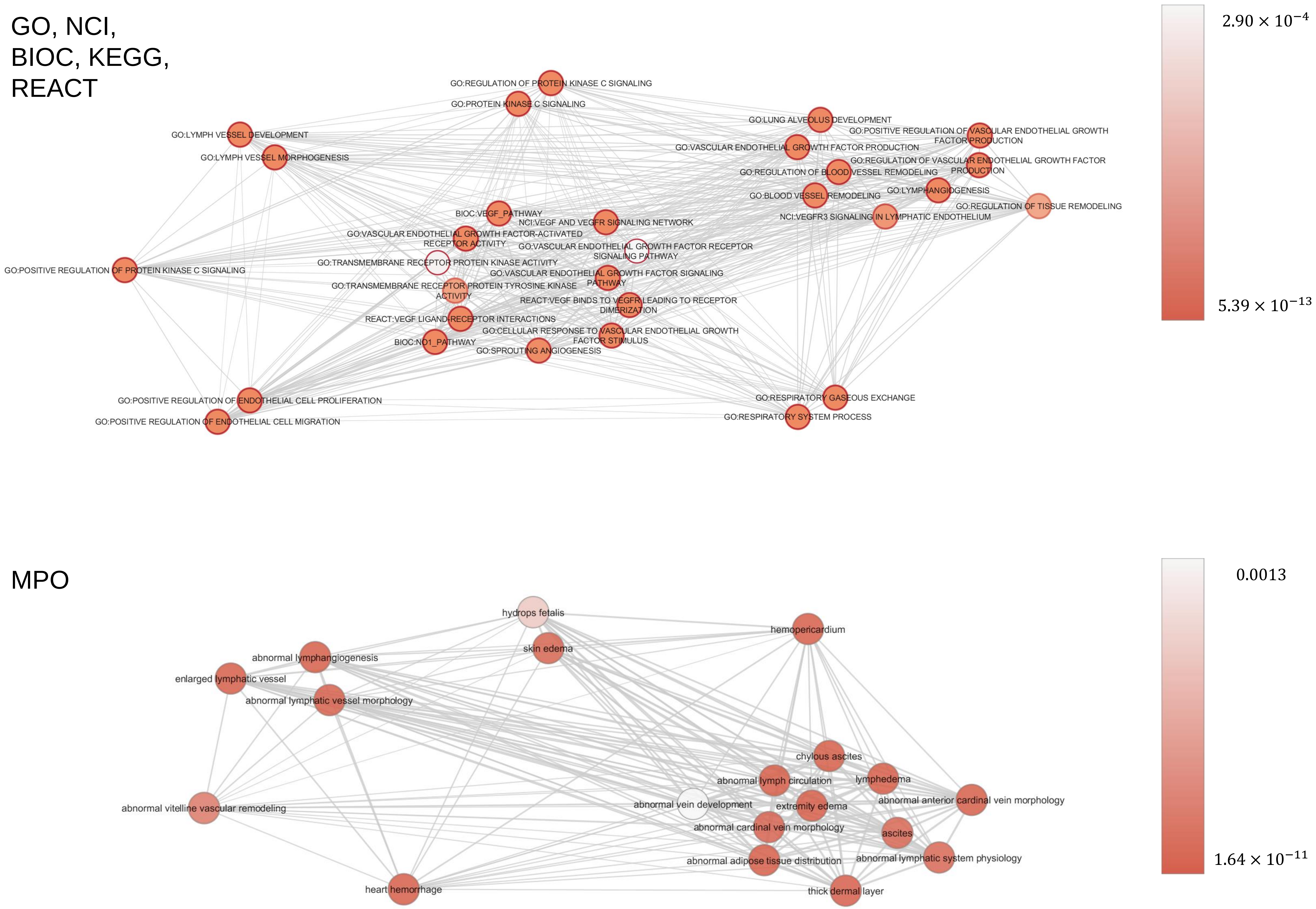
