## Supplementary Table S14) Patient phenotype and family history for selected deleterious missense and truncating variants for "Genes and pathways implicated in tetralogy of Fallot revealed by ultra-rare variant burden analysis in 231 genome sequences"

**Supplementary Table 14. Phenotype, family history and details of selected deleterious missense and truncating variants in 20 adults with TOF.**

| Case [a] | Phenotype and family history of CHD | Gene (transcript) | Variant [b] | Chromosomal position (GRCh37/hg19) |
|---|---|---|---|---|
| **Deleterious missense variants** | | | | |
| **TOF272** | TOF; VF and cardiac arrest age 39 y requiring ICD; hypothyroidism, died age 53 y; daughter with unspecified CHD | *NOTCH1* (NM_017617.3) | c.847T>G, p.(Cys283Gly) | chr9:139413913A>C |
| **TOF132** | TOF; endocarditis, depression, died age 59 y; infant daughter with TOF, persistent left SVC, solitary right thyroid lobe, Meckel's diverticulum, ectopic pancreas, died post-operatively | *NOTCH1* (NM_017617.3) | c.1243G>A, p.(Glu415Lys) | chr9:139412601C>T |
| **TOF310** | TOF; asthma; sister with unspecified CHD, died at 3 days | *NOTCH1* (NM_017617.3) | c.1816G>A, p.(Glu606Lys) | chr9:139410022C>T |
| **TOF220** [c] | TOF, RAA, APV; learning difficulties | *NOTCH1* (NM_017617.3) | c.1869C>A, p.(Asn623Lys) | chr9:139409969G>T |
| **TOF303** | TOF; depression | *NOTCH1* (NM_017617.3) | c.2045G>A, p.(Cys682Tyr) | chr9:139409124C>T |
| **TOF174** | TOF; Hageman factor XII deficiency | *NOTCH1* (NM_017617.3) | c.2128G>A, p.(Asp710Asn) | chr9:139409041C>T |
| **TOF57** | TOF, PA, RAA; anxiety, depression | *NOTCH1* (NM_017617.3) | c.2444G>A, p.(Cys815Tyr) | chr9:139407496C>T |
| **TOF131** | TOF, BAV; daughter with BAV and aortic coarctation, brother with hypoplastic left heart and aortic atresia, died at 9 days | *NOTCH1* (NM_017617.3) | c.4606T>C, p.(Cys1536Arg) | chr9:139399537A>G |
| **Truncating variants identified in the current study and not previously reported** | | | | |
| **TOF178** [d] | TOF, RAA, Late onset atrial flutter/fibrillation; depression | *FLT4* (NM_182925.4) | c.2766_2767dupCC, p.(Leu923Profs*4) | chr5:180046104dupGG |
| **TOF133** | TOF, PA, RAA; anxiety, depression, died age 49 y | *WNT5A* (NM_003392.4) | c.486C>A, p.(Cys162*) | chr3:55508563G>T |
| **TOF120** | TOF, PA, RAA; strabismus, ADHD, intussusception, scoliosis | *ZFAND5* (NM_001278245.1) | c.337_340delACTA, p.(Thr113Profs*118) | chr9:g.74974361delTAGT |

| Case [a] | Phenotype and family history of CHD | Gene (transcript) | Variant [b] | Chromosomal position (GRCh37/hg19) |
|---|---|---|---|---|
| **Truncating variants previously reported** | | | | |
| **TOF158** | TOF, RAA, paroxysmal atrial flutter requiring ablation, mild aortic dilatation; depression and/or anxiety, migraine, melanoma; daughter with TOF shown to have inherited the same *FLT4* variant | *FLT4* (NM_182925.4) | c.3574C>T, p.(Gln1192*) | chr5:180038443G>A |
| **TOF238** | TOF, RAA, MAPCA, PA; aortic dilatation | *FLT4* (NM_182925.4) | c.2499C>G, p.(Tyr833*) | chr5:180047216G>C |
| **TOF284** | TOF, MAPCA, inconclusive results about RAA; aortic valve replacement | *FLT4* (NM_182925.4) | c.1622dupG, p.(Gln542Profs*3) | chr5:180049766dupC |
| **TOF254** | TOF, APV; bilateral femoral vein occlusions; depression and/or anxiety | *FLT4* (NM_182925.4) | c.1172_1173delAG, p.(Glu391Glyfs*35) | chr5:180053196delCT |
| **TOF68** | TOF, RAA, APV; depression and/or anxiety | *FLT4* (NM_182925.4) | c.1037delC, p.(Thr346Argfs*7) | chr5:180055948delG |
| **TOF301** | TOF, RAA, paternal first cousin with suspected VSD | *FLT4* (NM_182925.4) | c.3331+1G>T, p.? | chr5:180041067C>A |
| **TOF109** | TOF, PFO or ASD, atrial flutter; obesity; mild cognitive and memory problems attributed to cerebral ischemia; brother died in infancy of suspected cyanotic CHD | *KDR* (NM_002253.2) | c.3287G>A, p.(Trp1096*) | chr4:55955875C>T |
| **TOF155** | TOF, PFO or ASD; depression and/or anxiety; gastroesophageal reflux | *KDR* (NM_002253.2) | c.2638C>T, p.(Arg880*) | chr4:55962486G>A |
| **TOF62** | TOF, RAA; learning difficulties | *FOXO1* (NM_002015.3) | c.580_586delGTGCCCT, p.(Val194Thrfs*137) | chr13:41239764delAGGGCAC |

All variants are heterozygous. Subjects in this table are of European descent, by study design. Obesity was defined as body mass index (BMI) consistently >30 as an adult. Short stature was defined as height <3rd percentile using standard adult growth curves (see Reuter et al., 2019).

[a] Case numbers of participants are the same as those used in a previous report of loss of function variants affecting the VEGF pathway in TOF (Reuter et al., 2019). Note that other high impact variants previously reported (Reuter et al., 2019) for genes *FLT4* (n=3) and *KDR* (n=2) do not appear here, if they were deleterious missense variants or in frame deletions, nor do variants in other VEGF genes that were not identified using the methods in the current study (see manuscript text for discussion).

[b] Underline indicates variants that alter evolutionarily conserved cysteine residues of *NOTCH1* (see Figure 3)

[c] Individual previously identified to have a structural variant - a multi-exon deletion within VEGF pathway gene *BCAR1* (Reuter et al., 2019).

[d] Identified on reanalysis using GATK3.7 (Nov 2018).

Abbreviations: APV, absent pulmonary valve; ASD, atrial septal defect; ADHD, attention deficit hyperactivity disorder; BAV, bicuspid aortic valve; CHD, congenital heart disease; ICD, implantable cardioverter defibrillator; MAPCA, major aortopulmonary collateral arteries; PA, pulmonary atresia; PFO, patent foramen ovale; RAA, right aortic arch; SVC, superior vena cava; TOF, tetralogy of Fallot; VF, ventricular fibrillation; VSD, ventricular septal defect.
